# Extraordinary Evasion of Neutralizing Antibody Response by Omicron XBB.1.5, CH.1.1 and CA.3.1 Variants

**DOI:** 10.1101/2023.01.16.524244

**Authors:** Panke Qu, Julia N. Faraone, John P. Evans, Yi-Min Zheng, Claire Carlin, Mirela Anghelina, Patrick Stevens, Soledad Fernandez, Daniel Jones, Ashish Panchal, Linda J. Saif, Eugene M. Oltz, Kai Xu, Richard J. Gumina, Shan-Lu Liu

## Abstract

Newly emerging Omicron subvariants continue to emerge around the world, presenting potential challenges to current vaccination strategies. This study investigates the extent of neutralizing antibody escape by new subvariants XBB.1.5, CH.1.1, and CA.3.1, as well as their impacts on spike protein biology. Our results demonstrated a nearly complete escape of these variants from neutralizing antibodies stimulated by three doses of mRNA vaccine, but neutralization was rescued by a bivalent booster. However, CH.1.1 and CA.3.1 variants were highly resistant to both monovalent and bivalent mRNA vaccinations. We also assessed neutralization by sera from individuals infected during the BA.4/5 wave of infection and observed similar trends of immune escape. In these cohorts, XBB.1.5 did not exhibit enhanced neutralization resistance over the recently dominant BQ.1.1 variant. Notably, the spike proteins of XBB.1.5, CH.1.1, and CA.3.1 all exhibited increased fusogenicity compared to BA.2, correlating with enhanced S processing. Overall, our results support the administration of new bivalent mRNA vaccines, especially in fighting against newly emerged Omicron subvariants, as well as the need for continued surveillance of Omicron subvariants.

## Introduction

Severe acute respiratory syndrome coronavirus 2 (SARS-CoV-2), the causative agent of the coronavirus disease 2019 (COVID-19) pandemic, continues to circulate across the globe while rapidly evolving. The beginning of 2022 was marked by the emergence of Omicron BA.1/BA1.1 variant, establishing a turning point in the pandemic with decreased pathogenicity^1–5^, increased transmissibility^2^, and enhanced immune escape^6–13^. During 2022, the prototype Omicron variant has given rise to numerous subvariants, with many displaying even higher extents of immune escape^9,14–22^, endangering the efficacy of vaccination efforts.

Following a few months of BA.5 dominance in the summer of 2022, a highly immune evasive^16,23,24^ Omicron subvariant, i.e., BQ.1.1, became the most prevalent in the United States; however, it is now being quickly supplanted by a new subvariant, XBB.1.5^25^. The XBB lineage was initially discovered in India in mid-August of 2022, resulting from a recombination event between two BA.2 lineages titled BA.2.10.1.1 and BA.2.75^26^. The emergence of this subvariant raised much alarm, as it has brought together a number of mutations in the spike (S) protein with established immune evasion functions, including R346T, G446S and F486S (**Fig. 1A**)^15^. Importantly, the efficacy of monoclonal antibody treatments^24^, both monovalent^23^ and bivalent^16,27^ vaccination strategies, as well as immunity stimulated by infection^23,27^, are all less effective against XBB. Recently, XBB has acquired two more mutations in the S protein, including G252V (XBB.1) and G252V+S486P (XBB.1.5) (**Fig. 1A**). The influence of these mutations on XBB.1 and XBB.1.5 is currently unknown, though mutations at residue F486, such as F486V, F486I, F486S, have been recurring among prior Omicron subvariants^26^, representing a critical evolutionary hotspot^28^. Given the rapid growth of XBB.1.5 in circulation in the United States and other parts of the world (**Fig. 1B** and **Fig. S1A**), it is crucial that we understand its impact on current public health measures.

**Figure 1:**
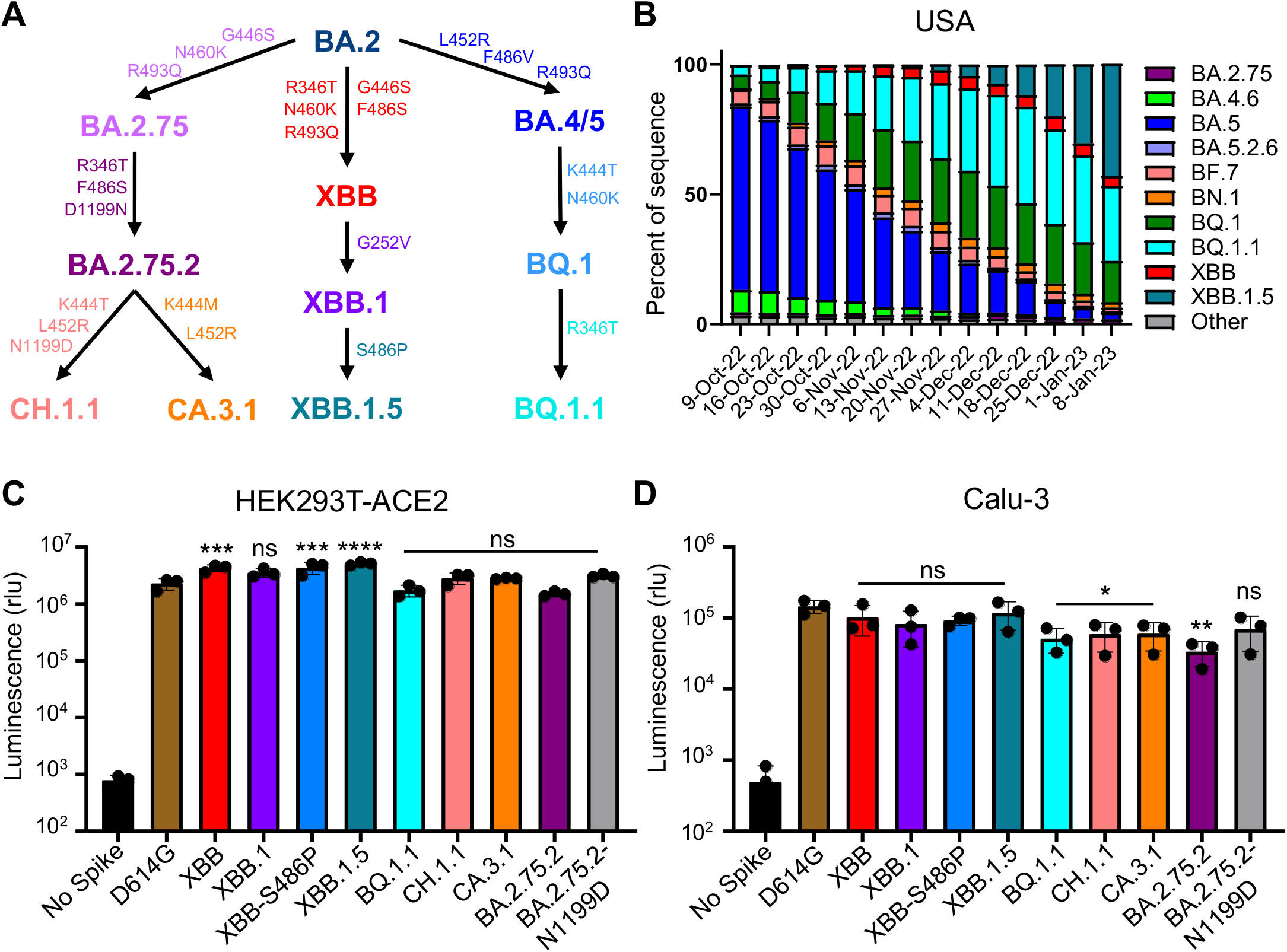
Distribution and infectivity of emerging Omicron subvariants XBB.1.5, CH.1.1, and CA.3.1. **(A)** Schematic depiction of the relationships between different Omicron subvariants, with key lineage-defining amino acid mutations for each displayed. **(B)** Distribution of recently emerged Omicron subvariants in the United States (USA) starting in early October of 2022 through the beginning of January 2023. Data were collected from the Centers for Disease Control and Prevention^25^ and plotted using Prism software. Infectivity of pseudotyped lentiviruses carrying each of the indicated S of the Omicron subvariants was determined in **(C)** HEK293T cells over expressing human ACE2 and **(D)** human lung cell-derived epithelial line CaLu-3. Transfection efficiency and S protein expression were comparable among all variants tested, as shown by western blotting in Figure 3E. Bars in **(B-C)** represent means ± standard deviation from three biological replicates. Significance relative to D614G was determined using a one-way repeated measures ANOVA with Bonferroni’s multiple testing correction (n=3). P values are displayed as ns p > 0.05, *p < 0.05, **p < 0.01, ***p < 0.001, and ****p < 0.0001.

In addition to BQ.1, BQ.1.1 and XBB subvariants, two other Omicron subvariants, CH.1.1 and CA.3.1, have also drawn attention. CH.1.1 emerged in Southeast Asia in November of 2022 and now accounts for more than 25% of infections in some parts of UK and New Zealand; it has caused alarm due to the appearance of the L452R mutation in the S protein^29^, which previously appeared in the more pathogenic Delta variant and highly transmissible BA.4/5 variants^18,30,31^. CA.3.1 emerged in the United States in December of 2022 and also carries this critical L452R mutation^29^. In this study, we investigate aspects of S protein biology of XBB.1.5, CH.1.1 and CA.3.1 in comparison to their parental variants, including entry into host cells, surface expression, fusogenicity, and processing. Most critically, we determine and compare their sensitivity to neutralizing antibodies stimulated by either bivalent or monovalent mRNA vaccination and previous infection (BA.5 wave), alongside with ancestral variants D614G, BA.2, and BA.2.75.2 as well as currently dominating variant BQ.1.1.

## Results

### Omicron subvariant XBB.1.5 exhibits an increase in viral infectivity, especially in CaLu-3 cells

First, we determined infectivity of lentiviruses pseudotyped with each of these subvariant S proteins in HEK293T cells stably expressing human ACE2 (HEK293T-ACE2) and human lung epithelial cell line CaLu-3. Both XBB and XBB.1.5 exhibited increased infectivity in HEK293T-ACE2 cells, with 1.9 times (p < 0.001) and 2.2 times (p < 0.0001) higher titer compared to D614G, respectively (**Fig. 1C**). The XBB.1.5 lineage-defining mutation S486P and G225V-containing XBB.1 also exhibited an increase in infectivity, with infectivity 1.9 times (p < 0.001) and 1.6 times (p > 0.05) higher than D614G, respectively (**Fig. 1C**). Of note, the infectivity of XBB.1.5 was not significantly higher than that of XBB (p > 0.05) (**Fig. 1C**). Interestingly, in contrast to the prototype Omicron BA.1 and subsequent subvariants that showed 3-5 times decreased infectivity in CaLu-3 cells^6,14,17^, the infectivity of these XBB subvariants was not significantly different from D614G, with titers only 1.4 times (p > 0.05) and 1.2 times (p > 0.05) lower than D614G for XBB and XBB.1.5, respectively (**Fig. 1D**). However, subvariants CH.1.1 and CA.3.1 still exhibited significant decreases in infectivity, with titers 2.5 times (p < 0.05) and 2.4 times (p < 0.05) lower than D614G, respectively (**Fig. 1D**). Consistent with previous results, BA.2.75.2 exhibited an infectivity 4.3 times lower than D614G, the lowest infectivity in CaLu-3 cells compared to D614G among all subvariants tested here (p < 0.01) (**Fig. 1D**). Together, these results appear to suggest that XBB.1.5, along with XBB, has gained an increased infectivity compared to the other Omicron subvariants although further investigation in primary lung epithelial cells and airway tissue is needed (see Discussion).

### Escape of neutralizing antibodies by XBB.1.5, CH.1.1, and CA.3.1 in bivalent vaccinated sera

In order to investigate neutralization resistance of emerging Omicron subvariants, we used our previously reported pseudotyped lentivirus neutralization assay^32^. We first examined their neutralization resistance to sera from 14 HCWs who had received a bivalent booster in addition to 2-4 doses of monovalent mRNA vaccine (n=14, 8 male and 6 female) (**Fig. 2A** and **Fig. 2D**). Among them, 12 HCWs received 3 doses of the monovalent Moderna mRNA-1273 or Pfizer BioNTech BNT162b2 vaccines followed by additional 1 dose of the bivalent Pfizer or Moderna vaccine, 1 HCW received 2 doses of the monovalent Pfizer BioNTech BNT162b2 vaccine and additional 1 dose of the bivalent Pfizer vaccine, and 1 HCW received 4 doses of the monovalent Pfizer BioNTech BNT162b2 vaccines plus additional 1 dose of the bivalent Pfizer vaccine (**Table S1**). Sera were collected between 23 and 108 days after receiving a bivalent vaccination (median 66).

**Figure 2:**
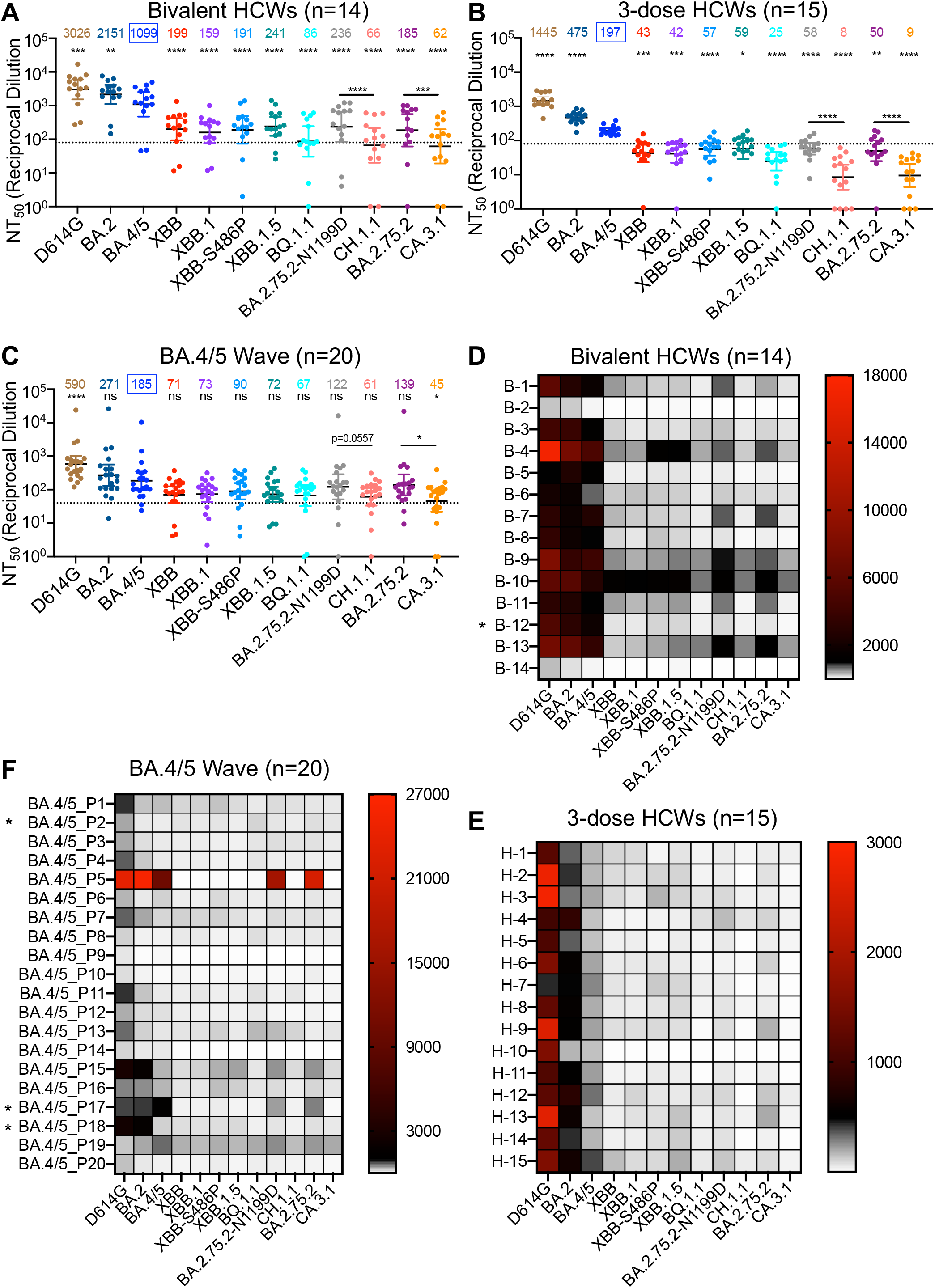
Neutralization of Omicron XBB.1.5, CH.1.1, and CA.3.1 subvariants by sera of bivalent or monovalent mRNA vaccinated health care workers (HCWs) and BA.4/5 wave infection. Neutralizing antibody titers were determined using lentiviral pseudotypes carrying each of the indicated S proteins of the Omicron subvariants. They were compared against BA.4/5 and/or respective parental Omicron subvariants as specified in the text. The cohorts included sera from 14 HCWs that received 3 monovalent doses of mRNA vaccine plus a dose of bivalent mRNA vaccine (n=14) **(A** and **D)**, 15 sera from HCWs that only received three doses of monovalent mRNA vaccine **(B** and **E)**, and 20 sera from BA.4/5-wave SARS-CoV-2 infected first responders and household contacts that tested positive during the BA.4/5 wave of infection in Columbus, Ohio **(C** and **F)**. Bars represent geometric means with 95% confidence intervals. Geometric mean NT_50_ values are displayed for each subvariant on the top. Statistical significance was determined using log_10_ transformed NT_50_ values to better approximate normality. Comparisons between multiple groups were made using a one-way ANOVA with Bonferroni post-test. And comparisons between two groups were performed using a paired, two-tailed Student’s t test with Welch’s correction. Dashed lines indicate the threshold of detection (80 for monovalent and bivalent mRNA vaccinees and 40 for BA.4/5 infection cohort. P values are displayed as ns p > 0.05, *p < 0.05, **p < 0.01, ***p < 0.001, ****p < 0.0001. Heatmaps in (**D-F**) depict neutralizing antibody titers by each individual against each Omicron subvariant tested. Asterisk in (**D**) indicates that the individual being infected by SARS-CoV-2 within six months before the sera sample collection, and asterisk in (**F**) indicates that the individuals who had received three doses of mRNA vaccines before infection.

Because of the continued dominance of Omicron subvariants, especially by BA.5 after the summer of 2022, all below comparisons for neutralization were made to BA.4/5 rather than the ancestral D614G. Strong neutralization resistance was exhibited by XBB.1.5, CH.1.1, and CA.3.1, with mean neutralizing antibody (nAb) titers 4.6 times (p < 0.0001), 16.7 times (p < 0.0001), and 17.7 times (p < 0.0001) lower than BA.4/5, respectively (**Fig. 2A** and **Fig. 2D**). Somewhat surprisingly, XBB.1.5 showed a modest increase in nAb titer compared to the parental XBB variant (**Fig. 2A** and **Fig. 2D**). However, neither of the two defining mutations for XBB.1.5 (G252V and S486P) contributed to the enhanced neutralization by bivalent sera, with nAb titers actually 6.9 times (p < 0.0001) and 5.8 times (p < 0.0001) lower than BA.4/5, respectively (**Fig. 2A** and **Fig. 2D**). Notably, BQ.1.1 exhibited a higher extent of neutralization resistance than all XBB subvariants, with nAb titer 12.8 times lower than BA.4/5 (p < 0.0001) and near the limit of detection (**Fig. 2A** and **Fig. 2D**). CH.1.1 and CA.3.1 are both derived from the subvariant BA.2.75.2, but the former has D1199N reversion mutation (**Fig. 1A**) denoted as BA.2.75.2-N1199D hereafter. We found that CH.1.1 had more immune escape than its parental BA.2.75.2-N1199D, with 2.7 times reduced nAb titer (p < 0.0001), whereas CA.3.1 exhibited stronger immune escape than its parental BA.2.75.2, with 3.0 times lower nAb titers (p < 0.0001) (**Fig. 2A** and **Fig. 2D**). Overall, we observed comparably strong immune escape among XBB subvariants that is less than BQ.1.1 but much more enhanced neutralization resistance for CH.1.1 and CA.3.1.

### XBB.1.5, CH.1.1, and CA.3.1 exhibit an almost complete escape of neutralizing antibodies in three-dose vaccinated sera

Next, we investigated neutralization resistance of these new Omicron subvariants in sera from Ohio State University Wexner Medical Center health care workers (HCWs) who had received 3 doses of monovalent mRNA vaccine (n=15) (**Fig. 2B** and **Fig. 2E**). Samples were collected 2-13 weeks after vaccination with a homologous booster dose of monovalent Pfizer/BioNTech BNT162b2 vaccine (n = 12) or Moderna mRNA-1273 (n = 3). These HCWs included 10 male and 5 female individuals and ranged from 26 to 61 years of age (median 33) (**Table S1**). The average nAb titers of 3-dose mRNA vaccine against D614G, BA.2 and BA.4/5 were about 2.1-5.6 times lower than those of bivalent mRNA vaccines (**Fig. A-B**); dramatic reductions in neutralization sensitivity were observed for XBB.1.5, CH.1.1 and CA.3.1, which exhibited complete escape from neutralizing antibodies, with mean nAb titers 3.3 times (p < 0.05), 24.6 times (p < 0.0001), and 21.9 times (p < 0.0001) lower than BA.4/5, respectively (**Fig. 2B** and **Fig. 2E**). Similarly, CH.1.1 and CA.3.1 subvariants also had dramatically lower nAb titers than their parental BA.2.75.2-N1199D and BA.2.75.2, respectively (**Fig. 2B** and **Fig. 2E**). Importantly, the overall trends for each subvariant in the 3-dose mRNA vaccine cohort remained similar to that of bivalent mRNA vaccination (**Fig. 2A-B** and **Fig. 2D-E**), and this was even more obvious for the subgroup (n=4) that had high nAbs titers (**Fig. S2A**).

### Omicron subvariants XBB.1.5, CH.1.1, and CA.3.1 are virtually resistant to neutralization by sera of BA.4/5 infection

We also examined neutralization resistance of XBB.1.5, CH.1.1, and CA.3.1 to sera from BA.4/5 infection wave among Columbus, Ohio first responders and household contacts that tested positive for COVID-19 (n=20) (**Fig. 2C** and **Fig. 2F**). Nasal swab samples were sequenced to identify the specific variant that caused infection, with 4 patients being infected by BA.4 or BA.4-derivative variants, 7 patients being infected with BA.5 or BA.5-derivative variants, and 9 patients being infected with undetermined SARS-CoV-2 variants (**Table S1**). All sample collection occurred during a BA.4 and BA.5 dominant period in Columbus, OH (July 2022 through September 2022). In this cohort, 17 individuals were unvaccinated, and 3 individuals had received 3 doses of either the Pfizer BioNTech BNT162b2 (n = 1) or Moderna mRNA-1272 (n = 2) vaccine (**Table S1**). Similar to the results shown above for the bivalent and monovalent mRNA vaccines, strong and almost complete neutralization resistance was observed for XBB.1.5, CH.1.1, and CA.3.1, with nAb titers 2.6 times (p > 0.05), 3.0 times (p > 0.05), 4.1 times (p < 0.05) lower than BA.4/5, respectively (**Fig. 2C, Fig. 2F** and **Fig. S2B**). Again, overall trends for each subvariants in this cohort followed the same patterns demonstrated in cohorts described for the bivalent and monovalent mRNA vaccines.

### Fusogenicity, surface expression, and processing of XBB.1.5, CH.1.1, and CA.3.1 S proteins

To investigate the biological function of the S proteins of these new Omicron subvariants, we investigated the fusogenicity, surface expression and processing. Consistent with our previous reports^15,17,33^, all subvariants tested exhibited reduced syncytia formation compared to the ancestral D614G (p < 0.0001), but with a clear increase in fusion relative to BA.2 (**Fig. 3A-B**). Like BQ.1.1 and BA.2.75.2^17^, subvariants XBB.1.5, CH.1.1 and CA.3.1 also showed enhanced fusogenicity compared to BA.4/5 (**Fig. 3A-B**). However, relative to the parental XBB, the XBB.1.5 subvariant and its two single mutants XBB.1 containing G252V and XBB-S486P did not demonstrate obvious differences in S fusogenicity (**Fig. 3A-B)**. The syncytia formation efficiency of CH.1.1 and CA.3.1 were comparable to that of BA.2.75.2-N1199D but seemed much lower than the parental BA.2.75.2 (**Fig. 3A-B**). The apparently higher fusogenicity of BA.2.75.2 as compared to BA.2 was consistent with our previous observations^17^. Importantly, differences in fusogenicity could not be attributed to differences in surface expression, as demonstrated by comparable levels of signal on cells expressing individual S proteins measured by flow cytometry (**Fig. 3C-D**).

**Figure 3:**
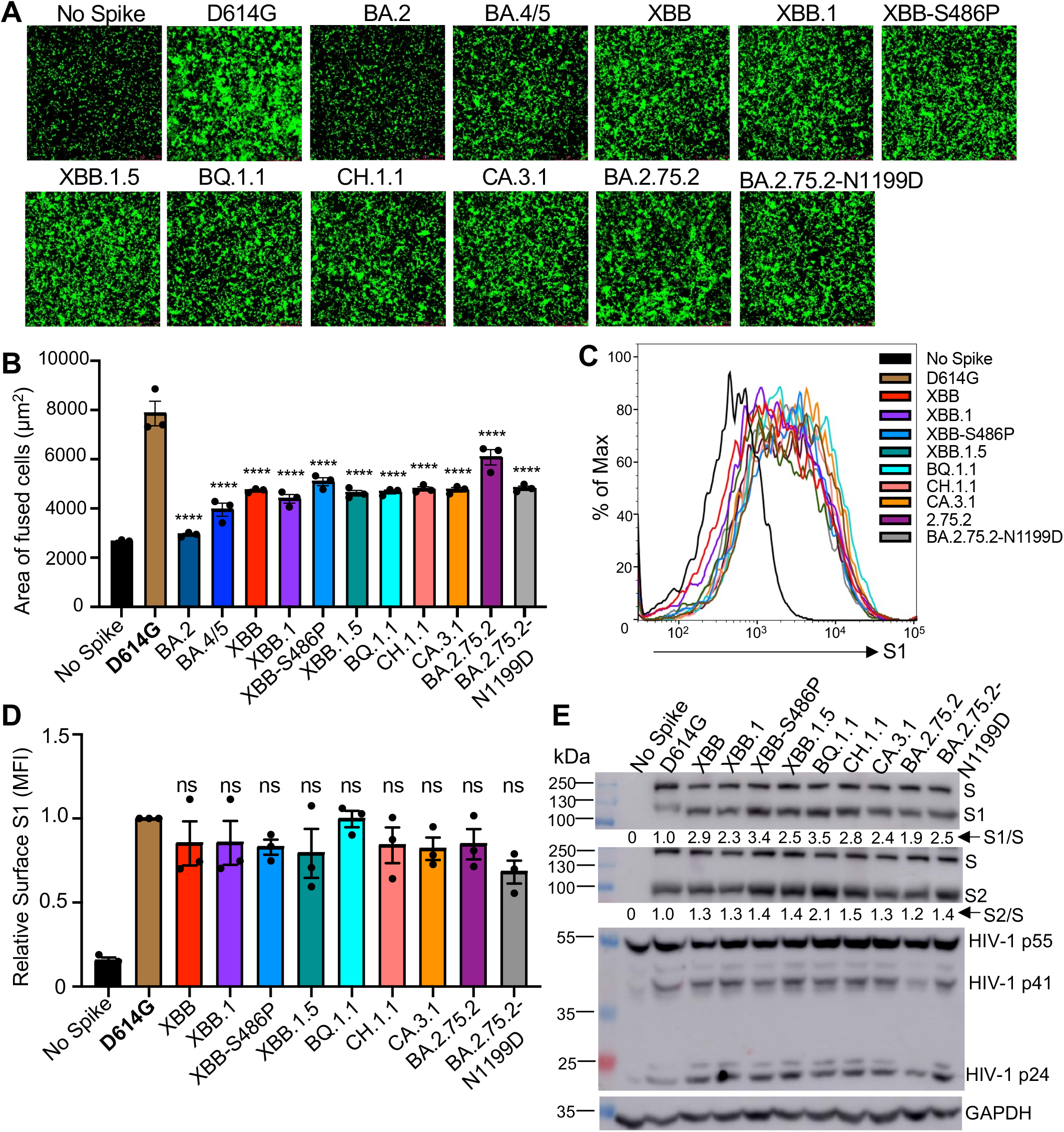
Syncytia formation, cell surface expression, and S processing of Omicron XBB.1.5, CH.1.1, and CA.3.1 subvariants. **(A-B)** Syncytia-forming activity. HEK293T-ACE2 cells were co-transfected with Omicron subvariant S proteins and GFP and incubated for 30 hours before **(A)** imaging and **(B)** quantifying syncytia. D614G and no S serves as positive and negative control, respectively. Comparisons in extents of syncytia for each variant were made against D614G, with p values indicating statistical significance. **(C-D)** Cell surface expression of S proteins. HEK293T cells used for production of pseudotyped lentiviral vectors bearing S proteins (Figures 1 and 2) from Omicron subvariants were fixed and surface stained for S with an anti-S1 specific antibody T62 followed by flow cytometric analyses. **(C)** Histogram plots of anti-S1 signals in transfected cells and **(D)** calculated relative mean fluorescence intensities of each subvariant by setting the value of D614G as 1.00. **(E)** S expression and processing. HEK293T cells used to produce pseudotyped vectors were lysed and probed with anti-S1, anti-S2, anti-GAPDH (loading control), or anti-p24 (HIV capsid, transfection control) antibodies; the signal for anti-S2 was from reblotting the membrane of anti-S1 and the signal for anti-GAPDH was from reblotting the membrane of anti-p24, respectively. S processing was quantified using NIH ImageJ and set to a S1/S or S2/S ratio and normalized to D614G (D614G = 1.0). Bars in **(B** and **D)** represent means ± standard error. Dots represent three biological replicates. Significance relative to D614G was determined using a one-way repeated measures ANOVA with Bonferroni’s multiple testing correction (n=3). P values are displayed as ns p > 0.05 and ****p < 0.0001.

Next, we investigated the S processing of these Omicron subvariants using pseudotyped viral producer cell lysates by immunoblotting. While the expression levels of these Omicron S proteins were comparable, all XBB subvariants, including XBB.1.5, CH.1.1 and CA.3.1, showed increased S processing compared to D614G; this was evidenced by increased S1/S and S2/S ratios, which was also true for BQ.1.1 and BA.2.75.2 consistent with our previous reports (**Fig. 3E**). Importantly, S processing for XBB.1.5 remained comparable to that for XBB, though a notable increase in S processing for XBB-S486P mutant was noticed (**Fig. 3E**), which was consistent with its relatively higher cell-cell fusion activity (**Fig. 3A-B**). No obvious differences in S processing for CH.1.1 and CA.3.1 were seen as compared to their parental BA.2.75.2 subvariant (**Fig. 3E**).

### Homology modeling reveals critical roles of lineage-defining mutations on XBB.1.5, CH.1.1 and CA.3.1 in receptor binding and immune evasion

To determine the impact of mutations S486P and G252V on immune evasion and receptor binding, we modeled the structures of XBB lineage spike protein in complex with either receptor ACE2 or representative neutralizing antibodies targeting these two residues. Located at the critical receptor binding domain (RBD)-ACE2 contact interface, residue F486 present in the parental BA2 subvariant is embedded in a hydrophobic groove formed by F28, L79, M82 and Y83 on ACE2; in contrast, the side chain of F486S or F486P, which are present in XBB/XBB.1 and XBB.1.5 subvariants, respectively, does not fit into the groove (**Fig. 4A**). In addition, comparing to the hydrophilic S486, residue P486 in XBB.1.5 is more hydrophobic, thus forming energetically favorable interactions with L79 and M82 on ACE2 and enabling better receptor utilization than XBB and XBB.1 (**Fig. 4A**). Moreover, residue 486 is an antigenic hotspot for class I neutralizing antibody recognition^28^. For example, therapeutic monoclonal antibody AZD8895 focuses its recognition on residue F486, with multiple antibody residues forming a surrounding hydrophobic cage; however, this interaction is abolished by the F486S/P mutations present in XBB and XBB.1.5 (**Fig. 4B**). Residue 252 is located on the spike N-terminal domain (NTD), which is also frequently recognized by neutralizing antibodies. **Fig. 4C** shows a representative NTD-targeting antibody (COVOX-159) which focuses its recognition on residue G252, yet a G252V mutation creates steric hindrance and abolishes this antibody recognition. Lastly, residues K444 and L452 are located within a common epitope site of class II RBD-targeting neutralizing antibodies (**Fig. 4D**); however, mutations in these two residues, i.e., K444T/M and L452R present in CH.1.1 and CA.3.1 subvariants, impact these interactions, leading to enhanced viral escape from established immunity induced by past vaccination or infection.

**Figure 4:**
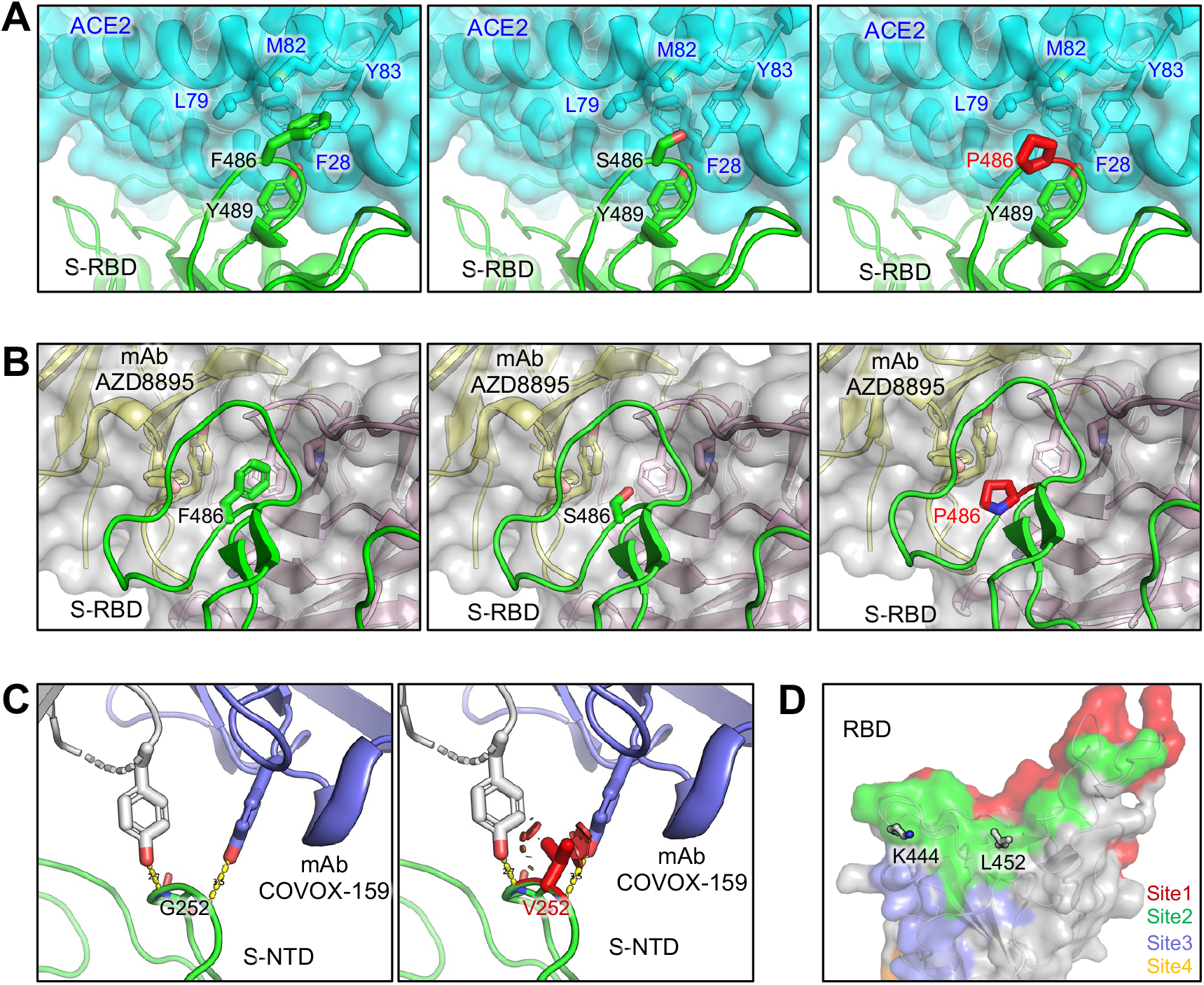
Homology modeling of key mutations in XBB.1.5, CH.1.1, and CA.3.1. (**A**) Structures of Spike receptor binding domain (RBD)-ACE2 binding interface shown as ribbons. (**B**) Structure of RBD with class I antibody AZD8895. The recognition focuses on residue F486, with multiple antibody residues forming a surrounding hydrophobic cage, whereas this interaction is abolished by F486S/P mutation. (**C**) Structures of an immune-dominant region of Spike N-terminal domain (NTD) with a representative antibody COVOX-159. The nAb recognition on residue G252 is abolished by G252V mutation through creating a steric hindrance (shown as red plates). (**D**) Residues K444 and L452 are located within a common epitope site of class II RBD-targeting neutralizing antibodies represented as green surface.

## Discussion

As SARS-CoV-2 continues to mutate and evolve, it is critical to monitor how the biology of the virus changes and the impact on the efficacy of current vaccines, including the currently bivalent mRNA vaccines. In this work, we found that the bivalent mRNA vaccine recipients exhibit approximately 2∼8-fold higher nAb titers, depending on variants tested, as compared to monovalent booster recipients, and the results are consistent with enhanced vaccine efficacy for the bivalent formula^34^. The nearly complete escape of 3-dose sera and BA.4/5 wave infection exhibited by all Omicron subvariants, especially XBB subvariants and CH.1.1 and CA.3.1, was remarkable and this is supported by some recent studies^23,24,27,35–37^. Notably, XBB.1.5 did not exhibit enhanced neutralization resistance over the recently dominant BQ.1.1 variant, which itself caused no notable surge in cases or hospitalizations compared to prior Omicron subvariants. Due to the fact that most samples fell below the limit of nAb detection, it is difficult to compare the neutralization titers between the different subvariants for this cohort. However, it is clear that CH.1.1 and CA.3.1 have a consistently stronger neutralization resistance than XBB, XBB.1 and XBB.1.5, which is astonishing and warrants continuous monitoring and further investigations.

One curious finding of this study is the modestly but consistently enhanced infectivity of Omicron XBB variants in CaLu-3 cells, especially XBB.1.5, as compared to the prototype Omicron BA.1/BA.1.1 and subsequently emerged Omicron subvariants including BQ.1.1 and BA.2.75.2 (**Fig. 1D**). We do not believe that these results are experimental artifacts, as assays were performed side by side at the same time for all variants and the expression levels of S proteins are also comparable for all variants. One possible explanation is the increased binding of XBB.1.5 to the ACE2 receptor, as recently shown by Richard Cao and colleagues^37^, which is also supported by our structural modeling. The initial mutation F486S in XBB is predicted to cause decreased affinity between the S protein and ACE2 due to the introduction of energetically unfavorable contacts between the polar residue and a hydrophobic patch. The subsequent mutation S486P largely reverses this effect, increasing the propensity for hydrophobic interactions with ACE2 and the flexibility of this region of the S protein, thus allowing it to settle further into the binding groove on ACE2 (**Fig. 4**). Consistent with the predicted improvement in ACE2 utilization, we observed a corresponding increase in cell-cell fusion and S processing for XBB subvariants, especially the single point mutant XBB-S486P (**Figs. 3A-B** and **3E**). Given all prior Omicron subvariants have been shown to exhibit low infectivity in CaLu-3 cells^6,9,14,17^, which correlated with their notably lower pathogenicity^1,4^ and a shift in tissue tropism toward the upper airway^2^, *in vivo* experiments investigating these aspects of the virus are necessary for XBB subvariants.

Our structural modeling of the spike protein interacting with its receptor and neutralizing antibodies provide insights for understanding Omicron subvariant evolution. Intriguingly, the structural analysis suggests a sophisticated two-step strategy for XBB lineage to evade immune suppression and likely outcompete other Omicron subvariants through mutations on the spike residue at position 486. F486 has a bulky hydrophobic side chain and is a hotspot for establishing protective immunity against the virus^17^ (**Fig. 1A**), while F486S mutation greatly facilitates evasion of antibody recognition. However, this F486S mutation reduces the efficiency of receptor utilization which must be counteracted with receptor-affinity gaining mutations, such as N460K and R493Q, to preserve the viral fitness. Thus, it makes sense that once the XBB circulation is established, S486 is further mutated into P486 as present in XBB.1.5, thus regaining higher receptor affinity while still maintaining similar immune escape. This combination of enhanced antibody escape and receptor affinity therefore likely enables, and has facilitated, the current dominance of the XBB.1.5 strain. In the cases of subvariants CH.1.1 and CA.3.1, it is clear that these variants have used the same strategy as other Omicron variants including BA.4/5 and BQ.1, by mutating the L452 and K444 sites of vulnerability frequently recognized by class I and II neutralizing antibodies to evade neutralization, again underscoring the convergent viral evolution.

Overall, our study highlights the continued waning of 3-dose mRNA booster efficacy against newly emerging Omicron subvariants. This effect can be partially saved by administration of a bivalent booster, though escape by some subvariants, particularly CH.1.1 and CA.3.1, is still prominent. Continued refinement of current vaccination strategies or investigation of new ones remains necessary. The biology of the S protein of Omicron subvariants, notably those of the XBB lineage, also continues to change, emphasizing the importance of continued surveillance of emerging variants.

### Limitations of Study

Throughout the study, cohorts of relatively small sample size are used to assess neutralizing antibody titers against the subvariants. However, previous studies have used cohorts of similar size and generated reliable data that has since been confirmed by other groups. Our cohorts also vary widely in time of sample collection after boosting or infection due to the clinical arrangements around collection of samples. In the bivalent cohort, 10 in 14 HCWs had been infected with SARS-CoV-2, with only one infected within 6 months of sample collection (Fig. 2D, B-12 denoted with *) and 9 infected more than 6 months before sample collection. Given our published results that mRNA booster vaccine-induced neutralizing antibody titer drops 17-20% every 30 days^38^, breakthrough infection in this bivalent cohort should not have had significant impact on their neutralization titers. A small subset of the BA.4/5 convalescent individuals (3 in 20) also received doses of vaccine, though we did not perform subgroup analysis due to the small size of the group. The use of pseudotyped virus instead of live virus for our assays is also a limitation, though we have previously validated our pseudotyped lentiviral system alongside live SARS-CoV-2^32^ and pseudotyped vectors are a common system used in the field. Finally, the influence of XBB.1.5, CH.1.1 and CA.3.1 signature mutations on ACE2 binding and antibody interaction warrants additional structural and biochemical characterization. Despite these limitations, the dramatic phenotypes of immune evasion by XBB subvariants and CH.1.1 and CA.3.1 are clear and have been corroborated by other studies^23,24,27,35–37^. Again, our study emphasizes the need for continued surveillance of emerging SARS-CoV-2 variants and the investigation of how viral evolution impacts vaccine efficacy and spike protein biology.

## Acknowledgements

We thank the NIH AIDS Reagent Program and BEI Resources for providing important reagents for this work. We also thank the Clinical Research Center/Center for Clinical Research Management of The Ohio State University Wexner Medical Center and The Ohio State University College of Medicine in Columbus, Ohio, specifically Francesca Madiai, Dina McGowan, Breona Edwards, Evan Long, and Trina Wemlinger, for logistics, collection, and processing of samples. In addition, we thank Sarah Karow, Madison So, Preston So, Daniela Farkas, and Finny Johns in the clinical trials team of The Ohio State University for sample collection and other supports. This work was supported by a fund provided by an anonymous private donor to OSU. S.-L.L., F.S., D. J., A.P., R.J.G., L.J.S. and E.M.O. were supported by the National Cancer Institute of the NIH under award no. U54CA260582. The content is solely the responsibility of the authors and does not necessarily represent the official views of the National Institutes of Health. J.P.E. was supported by Glenn Barber Fellowship from the Ohio State University College of Veterinary Medicine. R.J.G. was additionally supported by the Robert J. Anthony Fund for Cardiovascular Research and the JB Cardiovascular Research Fund, and L.J.S. was partially supported by NIH R01 HD095881.

## Author Contributions

S.-L.L. conceived and directed the project. R.J.G led the clinical study/experimental design and implementation. P.Q. performed most of the experiments. J.N.F. assisted in experiments. P.Q, J.N.F., and J.P.E. performed data processing and analyses. K.X. performed molecular modeling and data analyses. C.C., M.A., P.S., S.F., D.J., and R.J.G. provided clinical samples and related information. P.Q., J.N.F., J.P.E., and S.-L.L. wrote the paper. Y.-M.Z, L.J.S., E.M.O., and R.J.G. provided insightful discussion and revision of the manuscript.

---

**Figure S1:**
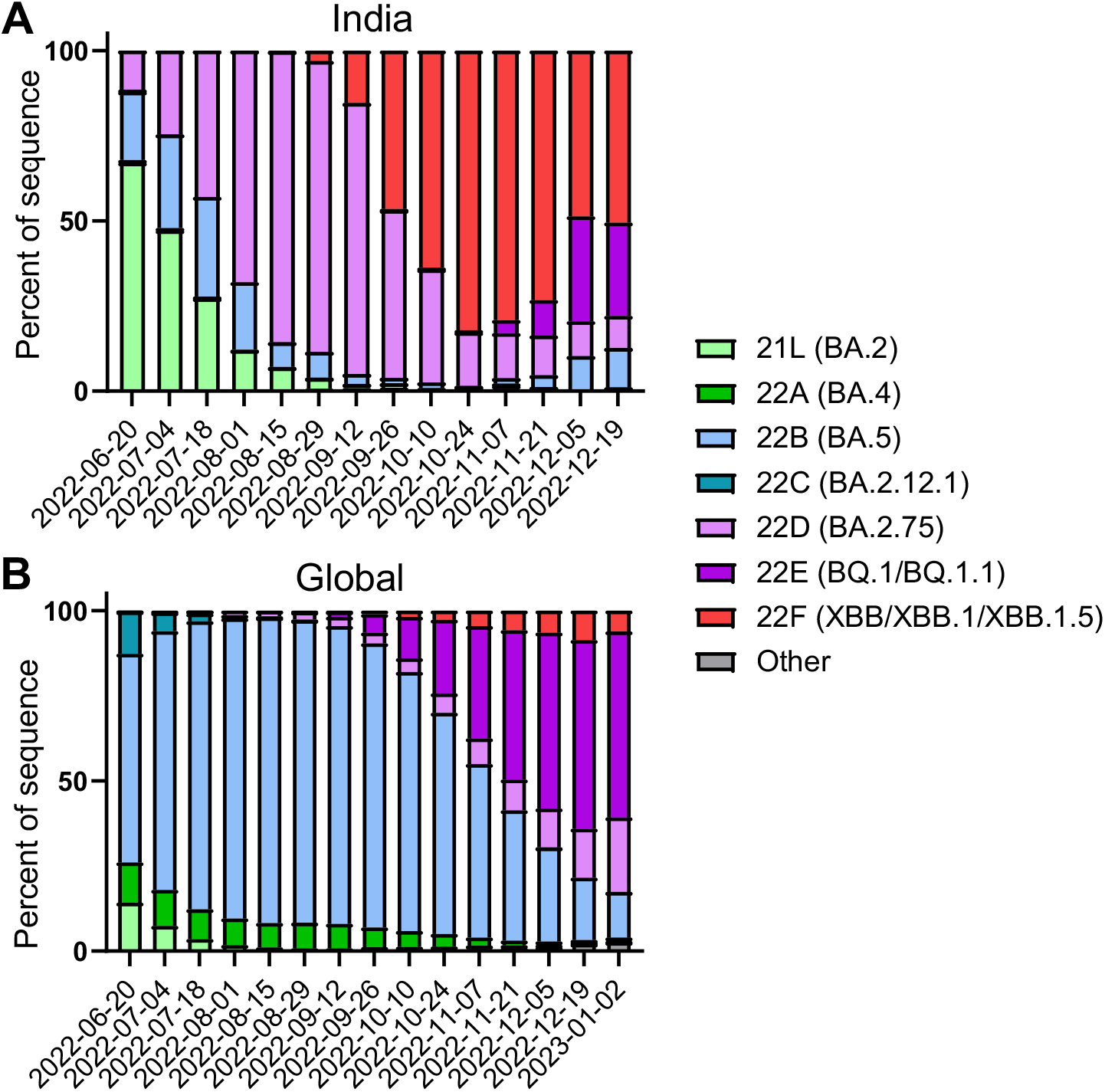
Distribution of recently emerged Omicron subvariants in India and around the world. (**A**) Distribution of indicated Omicron subvariants in India from June 20, 2022, to December 19, 2022. (**B**) Global distribution of indicated Omicron subvariants, collected staring from late June of 2022 through the beginning of January 2023. Data were collected from CoVariants^39^ and plotted using Prism software.

**Figure S2:**
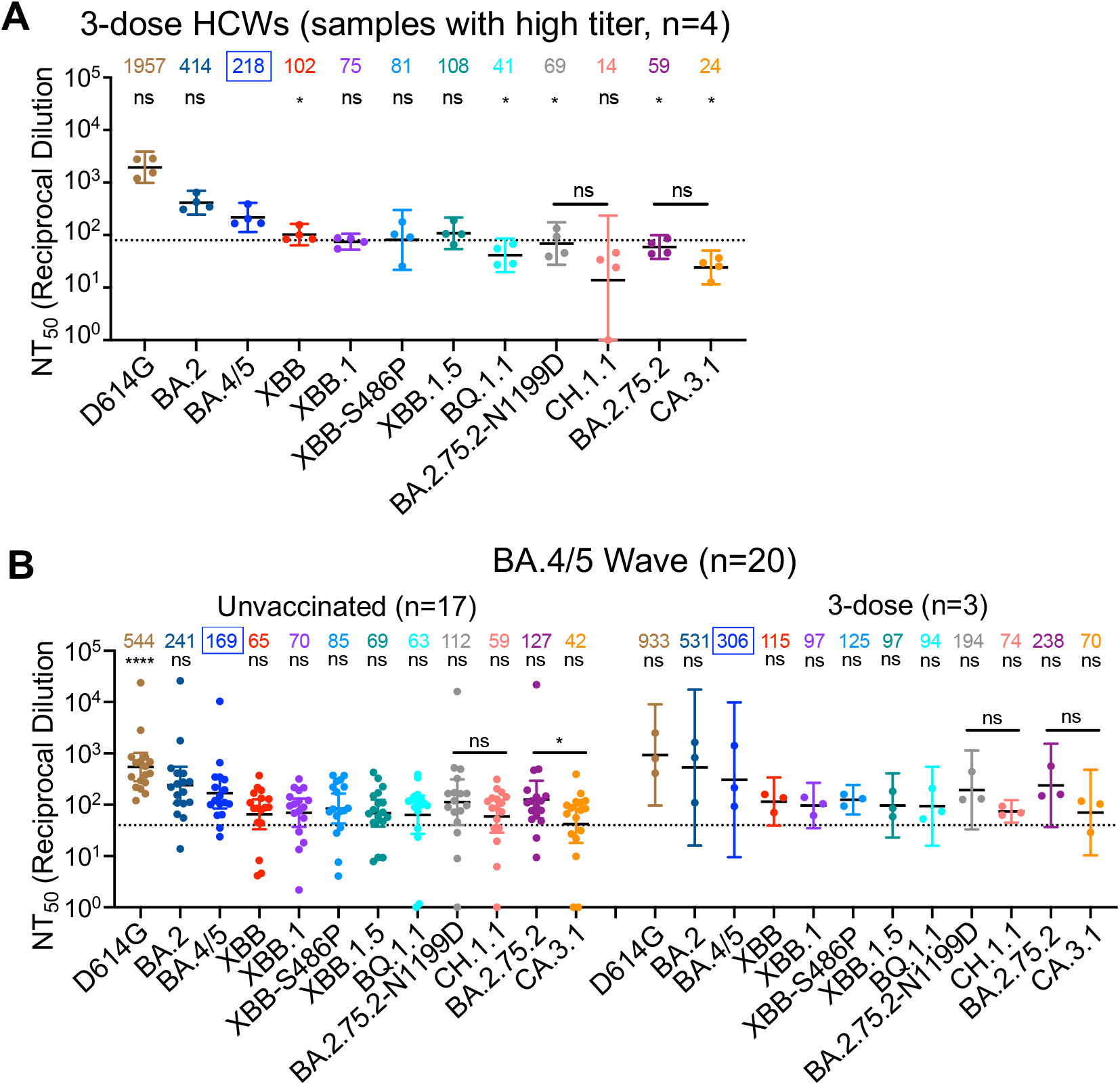
Subgroup analyses of neutralization. **(A)** Neutralization by 3-dose HCWs sera that exhibited high neutralizing antibody titers (n=4). Neutralizing antibody titers in 4 HCWs that received 3 homologous doses of monovalent mRNA vaccine were determined using pseudotyped lentiviral vectors carrying S proteins for each of the Omicron subvariants. Titers for the whole cohort are depicted in **(Fig. 2B)** while this figure focuses on samples that exhibited neutralizing antibody titers close to or above the limit of detection for XBB-derived variants (n=4). Bars represent geometric means with 95% confidence intervals. Geometric mean NT_50_ values are displayed for each subvariant. Dashed lines indicate the threshold of detection, i.e., 80. P values are displayed as ns p > 0.05 and *p < 0.05. (**B**) NT_50_ data shown in **Fig. 2C** for BA.4/5 sera is replotted by separating the vaccination status, i.e., 3-dose (n = 3) vs. unvaccinated (n = 17). Bars represent geometric means with 95% confidence intervals. Dashed lines indicate the threshold of detection, i.e., 40. Geometric mean NT_50_ values are displayed for each subvariant, with significance determined by one-way repeated measures ANOVA using Bonferroni’s multiple testing correction to make comparisons between multiple groups, and a paired, two-tailed Student’s t test with Welch’s correction to perform comparisons between two groups. P values are displayed as ns p > 0.05 and *p < 0.05.

## Materials and Methods

### Vaccinated and patient cohorts

Three cohorts were utilized in this study, the first being healthcare workers (HCWs) that received 3 homologous doses of mRNA vaccine. These samples were collected under approved IRB protocols 2020H0228, 2020H0527, and 2017H0292. The cohort included 15 HCWs that received homologous doses of either the monovalent Moderna mRNA-1273 (n=3) or the monovalent Pfizer BioNTech BNT162b2 (n=12) vaccines. Samples were collected from 14 to 86 days post-booster vaccination (median 40). HCW ages ranged from 26 to 61 (median 33). The cohort included 10 male and 5 female individuals.

The second cohort were HCWs that received a bivalent mRNA booster formulation. These samples were collected following the approved IRB protocols 2020H0228, 2020H0527, and 2017H0292. The cohort included 1 HCW that received 2 doses of the monovalent Pfizer BioNTech BNT162b2 vaccines and additional 1 dose of the bivalent Pfizer vaccine, 12 HCWs that received 3 doses of the monovalent Moderna mRNA-1273 or Pfizer BioNTech BNT162b2 vaccines and additional 1 dose of the bivalent Pfizer or Moderna vaccine, and 1 HCW that 4 doses of the monovalent Pfizer BioNTech BNT162b2 vaccines and additional 1 dose of the bivalent Pfizer vaccine. Samples were collected from 23 to 108 days post-bivalent vaccination (median 66). HCW ages ranged from 25 to 48 (median 36). The cohort included 8 male and 6 female individuals.

The third cohort included first responders and their household contacts that tested positive for SARS-CoV-2 infection during the BA.4/5 wave in Columbus, OH. These samples were collected under approved IRB protocols 2020H0527, 2020H0531, and 2020H0240. The cohort included 20 individuals. For each individual, a nasal swab was collected and sequenced to confirm the variant they were infected with. 4 individuals were infected with BA.4 and 7 individuals were infected with BA.5. The infecting variant could not be determined for the remaining 9 individuals but the dates of collection fall within when BA.4/5 was dominant in Columbus, OH (late July 2022 through late September 2022). Ages ranged from 27 to 58 years (median 44) and the cohort included 4 male and 15 female individuals. The age and gender of one individual are unknown. The cohort included 17 individuals that were unvaccinated and 3 individuals that received 3 homologous does of either the monovalent Pfizer BioNTech BNT162b2 (n=1) or the monovalent Moderna mRNA-1273 (n=2) vaccines.

### Cell lines and maintenance

Human embryonic kidney cell lines HEK293T (ATCC CRL-11268, RRID: CVCL_1926) and HEK293T engineered to overexpress human ACE2 (BEI NR-52511, RRID: CVCL_A7UK) were maintained in DMEM (Gibco, 11965-092) supplemented with 10% FBS (Sigma, F1051) and 0.5% penicillin-streptomycin (HyClone, SV30010). Human adenocarcinoma lung epithelial cell line CaLu-3 (RRID: CVCL_0609) was maintained in EMEM (ATCC, 30-2003) supplemented with 10% FBS and 0.5% penicillin-streptomycin. The cells were incubated at 37 °C and 5.0% CO_2_. Passaging of all cell lines was performed by first washing with Dulbecco’s phosphate buffer saline (Sigma, D5652-10×1L) followed by an incubation in 0.05% Trypsin + 0.53 mM EDTA (Corning, 25-052-CI) until complete cell detachment for splitting.

### Plasmids

Pseudotyped lentiviral vectors were produced as previously described^32^. Briefly, vectors are produced through the co-transfection of the HIV-1 vector pNL4-3 with an Env deletion and the SARS-CoV-2 spike of interest. The pNL4-3 vector includes a *Gaussia* luciferase reporter gene that is secreted by target cells. SARS-CoV-2 spike plasmids were generated in the pcDNA3.1 plasmid backbone either through KpnI and BamHI restriction enzyme cloning by GenScript Biotech (Piscataway, NJ) (D614G, BA.2, BA.4/5, and XBB) or site-directed mutagenesis via PCR (XBB.1, XBB-S486P, XBB.1.5, BQ.1.1, CH.1.1, CA.3.1, BA.2.75.2, and BA.2.72.2-N1199D) and confirmed by Sanger sequencing. All spike constructs include N- and C-terminal FLAG tags.

### Pseudotyped lentivirus production and infectivity

Pseudotyped lentiviral vectors were produced as previously described^32^. Briefly, HEK293T cells were co-transfected in a 2:1 ration with the pNL4-3-inGluc vector and the spike plasmid of interest using polyethyleneimine transfection (Transporter 5 Transfection Reagent, Polysciences) to produce pseudotyped lentiviral particles. The lentivirus was collected by taking the media of the transfected cells 48 and 72 hours post-transfection. Relative infectivity of the lentivirus was then assessed in both HEK293T-ACE2 and CaLu-3 cells. *Gaussia* luciferase activity measured at 72 hours post infection for HEK293T and 72 hours for CaLu-3 were used to determine relative infectivity. *Gaussia* luciferase activity was determined by taking equal volumes of infected cell media and *Gaussia* luciferase substrate (0.1 M Tris pH 7.4, 0.3 M sodium ascorbate, 10 µM coelenterazine) and combining for an immediate luminescence signal detected by a BioTek Cytation plate reader.

### Virus neutralization assay

Pseudotyped lentiviral neutralization assays were performed as previously described^32^. First, all serum samples were diluted 4-fold (final dilutions 1:80, 1:320, 1:1280, 1:5120, 1:20480, and no serum control for HCWs samples; final dilutions 1:40, 1:160, 1:640, 1;2560. 1:10240, and no serum control for BA.4/5-Wave samples). An equal volume of pseudotyped lentivirus was then added to the diluted sera and incubated at 37 °C for 1 hour. This neutralized virus mixture was then used to infect HEK292T-ACE2 cells. *Gaussia* luficerase activity was then determined 48 and 72 hours post infection. 50% neutralization titers (NT_50_) were determined by least-squares-fit, non-linear regression in GraphPad Prism 9 (San Diego, CA).

### Syncytia formation

To measure the extent of cell-cell fusion mediated by the different SARS-CoV-2 spikes, HEK293T cells expressing ACE2 were co-transfected with spike plasmid and GFP^6^. Cells were imaged with a Leica DMi8 confocal microscope 30-hours post-transfection. Representative images were selected for presentation. Area of fused cells was determined and quantified using the Leica X Applications Suite, scale bars represent 150 µM.

### S protein surface expression

HEK293T cells used to produce lentiviral vectors were harvested 72-hours post-transfection. Cells were incubated in PBS+5mM EDTA for 7 minutes at 37°C to mediate disassociation. The cells were fixed in 3.7% formaldehyde and stained with anti-SARS-CoV-2 polyclonal S1 antibody (Sino Biological, 40591-T62; RRID: AB_2893171) and secondary antibody anti-Rabbit-IgG-FITC (Sigma, F9887, RRID: AB_259816). S surface expression was measured using a Life Technologies Attune NxT flow cytometer and data was processed using FlowJo v7.6.5 (Ashland, OR).

### S protein processing

Lysate from HEK293T cells used to produce lentiviral vectors was collected through a 40-minute incubation on ice in RIPA lysis buffer (50mM Tris pH 7.5, 150 mM NaCl, 1 mM EDTA, Nonidet P-40, 0.1% SDS) supplemented with protease inhibitor (Sigma, P8340). Samples were run on a 10% acrylamide SDS-PAGE gel and transferred to a PVDF membrane. Membranes were probed with anti-S1 (Sino Biological, 40591-T62; RRID:AB_2893171), anti-S2 (Sino Biological, 40590; RRID:AB_2857932), anti-p24 (NIH HIV Reagent Program, ARP-1513), and anti-GAPDH (Santa Cruz, Cat# sc-47724, RRID: AB_627678). Secondary antibodies included Anti-Rabbit-IgG-HRP (Sigma, A9169; RRID:AB_258434) and Anti-Mouse (Sigma, Cat# A5278, RRID: AB_258232). Blots were imaged using Immobilon Crescendo Western HRP substrate (Millipore, WBLUR0500) and exposed on a GE Amersham Imager 600. Band intesnsities were quantified using NIH Image J analysis software (Bethesda, MD).

### Structural modeling and analyses

Structural modeling of XBB spike proteins in complex with either ACE2 receptor or neutralizing antibodies was performed on SWISS-MODEL server using published X-ray crystallography or cryo-EM structures as templates (PDB IDs: 7K8Z, 8DT3, 7L7D, 7XB0, 7NDD). Molecular contacts of XBB mutants were examined and illustrated with PyMOL.

### Quantification and statistical analysis

All statistical analyses were performed using GraphPad Prism 9 and are described in the figure legends. NT_50_ values were determined by least-squares fit non-linear regression in GraphPad Prism 9. Error bars in **(Fig. 1C-D)** represent means ± standard deviation and in (**Fig. 3B** and **Fig. 3D)** represent means ± standard error. Error bars in **(Fig. 2A-C** and **Fig. S2)** represent geometric means with 95% confidence intervals. Statistical significance was determined using log10 transformed NT_50_ values to better approximate normality (**Fig. 2A-C**, and **Fig. S2A-B**), comparisons between multiple groups were made using a one-way ANOVA with Bonferroni post-test, and a paired, two-tailed Student’s t test with Welch’s correction was used (**Fig. 2A–C** and **Fig. S2A-B**).

### Data and code availability

This paper does not report original code. NT_50_ values and de-identified patient information can be shared by the lead contact upon request. Any other additional data can be provided for reanalysis if requested from the lead contact.

---

**Table S1:** Demographic and sample collection information of HCWs and BA.4/5-Wave first responders and household contacts. Summary information for HCW sera samples collected 1-dose Pfizer or Moderna bivalent mRNA vaccine or samples collected 3-dose of mRNA vaccines is shown. Additionally, summary information of the BA.4/5-Wave Columbus, Ohio first responders and household contacts is also provided. na means “not applicable”.

|  | Bivalent HCWs<br>(n=14) | 3-dose HCWs<br>(n=15) | BA.4/5-Wave first<br>responders and<br>household contacts<br>(n=20) |
| --- | --- | --- | --- |
| **Age in Years at Sample Collection**<br>[Median (Range)] | 36 (25-48) | 33 (26-61) | 44 (27-58) |
| **Gender** [n (% of Total)] |  |  |  |
| Male | 8 (57.1%) | 10 (66.7%) | 4 (20%) |
| Female | 6 (42.9%) | 5 (33.3%) | 15 (75%) |
| Unknown | na | na | 1 (5%) |
| **Sample Collection Window** | Dec.<br>2022-early<br>Jan. 2023 | Aug. 2021-Feb. 2022 | July. 2022-Sep. 2022 |
| **Vaccine status** [n (% of Total)] |  |  |  |
| Unvaccinated | na | na | 17 (85%) |
| 3-dose Moderna | na | 3 (20%) | 2 (10%) |
| 3-dose Pfizer | na | 12 (80%) | 1 (5%) |
| 2-dose Pfizer+1-dose Pfizer bivalent | 1 (7.1%) | na | na |
| 3-dose Pfizer/Moderna+1-dose Pfizer/Moderna<br>bivalent | 12<br>(85.8%) | na | na |
| 4-dose Pfizer+1-dose Pfizer bivalent | 1 (7.1%) | na | na |
| **Sample Collection Timing** [Median (Range)] |  |  |  |
| Days post 3<sup>rd</sup> dose for Recipients of three doses | na | 40 (14-86) | 252 (141-279) |
| Days post the bivalent dose for Recipients | 66 (23-108) | na | na |
| **Infected Variants** |  |  |  |
| BA.4 or BA.4-derivative variants | na | na | 4 (20%) |
| BA.5 or BA.5-derivative variants | na | na | 7 (35%) |
| Undermined | na | na | 9 (45%) |

